# Unveiling Gene Regulatory Network Dynamics using Fuzzy Clustering

**DOI:** 10.64898/2025.12.31.697266

**Authors:** Rafael Kollyfas, Marta Cagna, Alexandra M. Nicaise, Ludovic Vallier, Irina Mohorianu

**Author notes:** Corresponding author: I Mohorianu.

## Abstract

Partitioning cells into robust, reproducible clusters is a core step across single-cell-resolution analyses; current state-of-the-art approaches struggle with capturing and summarising dynamics on continuous expression patterns. We present *Flufftail* (**F**uzzy **L**ogic **U**nifying **F**ramework reveals **T**ranscriptional **A**rchitectures summarised via **I**ntegrated **L**earning), an R framework and interactive Shiny app that consolidates clustering uncertainty by aggregating iterative stochastic partitions, resulting from a constant input. *Flufftail* computes per-cell membership probabilities, element-centric consistency scores, consensus matrices, and collapsed hard/crisp cell-assignments; we also leverage fuzzy assignments to prioritise genes that might act as regulatory hubs, subsequently using these as anchor points to infer gene regulatory network dynamics across transitions. We showcase the approach on single-nuclei and spatial transcriptomic case studies, illustrating how fuzzy clustering highlights transitional cell populations, proposing an ordered, state-dependent rewiring of regulatory interactions directly linked to the observed phenotype.

## Introduction

High-throughput sequencing technologies reshaped the study of biological systems by enabling genome-wide interrogation of gene expression and an in-depth assessment of regulatory interactions, leading to mechanistic hypotheses on gene-gene interplay. Early approaches focused on static gene regulatory networks (GRNs)[1, 2, 3]. The rapid expansion of single-cell and spatial (transcriptomic) technologies shifted the focus from sample-wide averages to gradients of expression mapped on cellular heterogeneity and spatial context [4]. The technical characteristics of these newer datasets also prompted a reassessment of the robustness and reproducibility of outputs obtained using canonical machine learning and data science approaches.

Clustering single-cell RNAseq (scRNAseq) datasets using community detection algorithms, e.g. Louvain [5] and Leiden [6], represents a fundamental initial step in current transcriptomics analyses. However, these algorithms are inherently stochastic, i.e. repeated runs on the same input can yield different clustering solutions in part due to the initialisation of the random seed. This stochasticity introduces variability in interpretation, with direct consequences for downstream analyses. Standard analysis pipelines such as Seurat [7] and Monocle [8] typically rely on a single clustering solution, implicitly assuming that it captures the underlying biological structure. By overlooking clustering variability, rare or transitional cell states, may be missed. Such cells often show unstable, yet informative, stable and reproducible assignments across repeated runs [9], reflecting genuine biological plasticity [10] rather than technical noise [11].

The aim of robust, reliable clustering is to recover true biological structure, while minimizing sensitivity to technical artefacts, e.g. random noise [11] or sequencing bias/stochasticity. Traditional cluster-centric metrics such as the Rand Index [12], focus on overall partition similarity neglecting the assessment of instability at individual-cell level. Element-centric consistency (ECC), also referred to as frustration [13], quantifies how consistently each point (in this context, cell) is assigned across repeated clustering runs [9]. This cell-centric perspective enables the identification of cells with reproducible yet inconsistent/fuzzy assignments, which may reflect biologically meaningful heterogeneity. For an iterative set of partitions, a high median ECC indicates that the clustering faithfully captures biological signal, rather than algorithmic artefacts. However, the assumption that a cell uniquely belongs to a single, stable cluster often fails in biological systems, where transitional or multi-phenotypic states are common [14]. For instance, Gribben et al. [10] reported the emergence of bi-phenotypic hepatocyte–cholangiocyte cells during late-stage chronic liver disease through iterative subclustering complemented by expert curation of unstable cells.

To systematically capture and characterize probabilistic cell assignment in single-cell-resolution clustering, we present *Flufftail* : (**F**uzzy **L**ogic **U**nifying **F**ramework reveals **T**ranscriptional **A**rchitectures summarised via **I**ntegrated **L**earning), a robust and scalable R package, accompanied by an interactive shiny interface. By aggregating results across multiple random seeds, *Flufftail* captures clustering variability on per-cell membership probabilities, ECC scores, robust hard cluster assignments via majority voting, and a consensus matrix capturing co-clustering behaviour. These outputs enable the identification and characterization of ambiguous cells with unstable cluster assignments, which are often enriched for transitional states. Our central hypothesis is that residual instability in a small subset of cells, reflected by low ECC within an otherwise stable (high median ECC) clustering, reflects biologically interesting plasticity rather than technical noise [15]. Existing approaches relying on consensus clustering, such as SC3 [16] and scALPO [17]), quantify uncertainty at cluster level but overlook the downstream regulatory consequences of cellular fuzziness. This gap is critical, as emerging evidence highlights cellular plasticity as a driver of dynamic behaviour and variability in therapeutic response [10, 18].

## Materials and Methods

### Datasets

We developed and evaluated *Flufftail* on real and synthetic single-cell and spatial transcriptomics datasets, to thoroughly assess robustness, biological relevance, and scalability. The primary dataset used in this study was the single-nuclei RNAseq dataset from Gribben et al. [10], focusing on hepatocytes and cholangiocytes from end-stage liver disease samples. This dataset was selected to show case how *Flufftail* captures cells transitioning between these cell types, recently described as bi-phenotypic cells [10]. After removing a small, spatially distinct hepatocyte island, the Seurat object [7] comprised 23,835 nuclei across 29,653 genes; the expression matrix was preprocessed with SCTransform [19], reduced by PCA, and visualised using UMAP [20].

To emphasise the generalisability of *Flufftail* to spatial assays, we applied the framework to the Alsema et al. [21] multiple sclerosis spatial transcriptomics dataset, restricting analyses to tissue section A1 and combining the top 500 highly variable genes with curated gene sets to highlight niche-relevant programmes.

To benchmark the scalability of *Flufftail*, we simulated datasets with Splatter [22] up to 100,000 cells. Performance was evaluated on a 48-core server with 754 GB RAM and on a MacBook Pro (Apple M3 Pro, 18 GB RAM), reporting runtime and memory usage across both environments.

All analyses were performed in R v4.2 (seed = 42). The code, with example case studies, is provided in the *Flufftail* GitHub repository to enable full reproducibility (https://github.com/Core-Bioinformatics/Flufftail).

### *Flufftail* Overview

*Flufftail* is a modular framework (Figure 1) designed to capture clustering uncertainty in single-cell-resolution datasets and to link this variability to downstream analyses modelling cellular transitions and regulatory dynamics. The workflow is organised into a series of stages, each described in detail in the following subsections.

**Figure 1.**
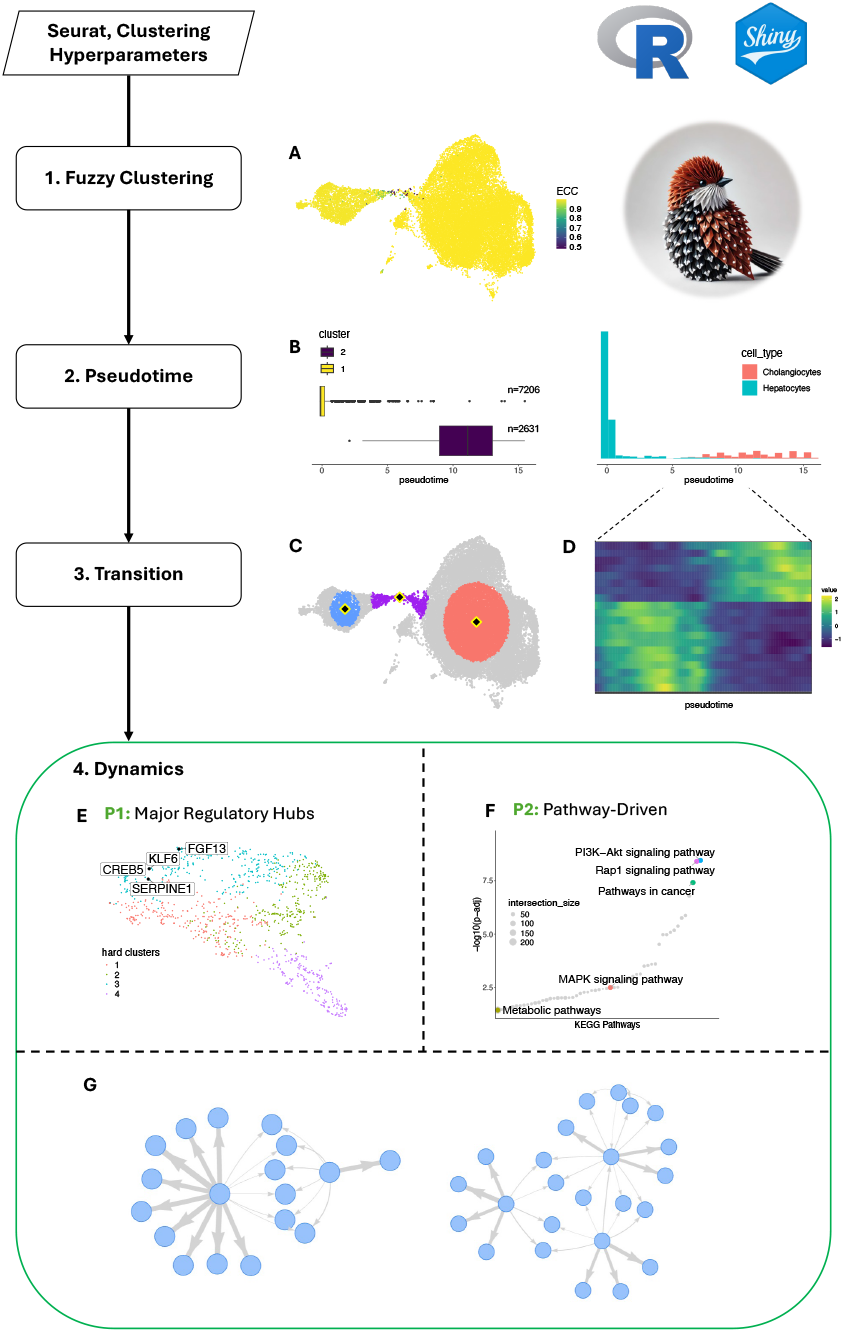
Overview of the *Flufftail* pipeline, illustrated on a case study highlighting regulatory dynamics at single-cell resolution. Key analytical steps comprise repeated stochastic clustering, identification of fuzzy cells, prioritisation of major regulatory hubs, and inference of gene regulatory network (GRN) dynamics. **(A)** ECC-weighted UMAP, following repeated stochastic clustering; the cells with lowest ECC scores (green/purple gradient) are localised on a bridge between the two main cells types, illustrating cell-state plasticity, as bi-phenotypic cells (*Gribben et al*). **(B)** Decomposed pseudotime showing the distribution of annotated cell types relative to computationally identified clusters along the inferred Monocle pseudotime trajectory (x-axis). **(C)** UMAP highlighting cells sampled around the centroids of two stable clusters and their midpoint (i.e. unstable cells), defining the main transition zone used for downstream analyses. **(D)** Heatmap of expression gradients across pseudotime (x-axis) for the top 20 differentially expressed genes ordered along the low ECC transition. **(E)** Gene-module UMAP highlighting four *Flufftail*-identified major regulatory hubs (KLF6, SERPINE1, CREB5, and FGF13), transcriptionally upregulated in biphenotypic hepatocyte–cholangiocyte cells (KLF6 and SERPINE1 were experimentally validated as core plasticity markers, *Gribben et al*). **(F)** Pathway-driven approach highlighting dynamic genes resulting from a gene set enrichment analysis. **(G)** Show-case panel illustrating the concept of GRN dynamics, focusing on changes in topology (rewiring events) and changes in interaction strength (proportional to the edge width).

Repeated stochastic clustering is used to quantify the variability in cell assignments; next, we derive fuzzy and consensus representations of cluster structures (see *Fuzzy and Consensus Clustering*). Cells with unstable/fuzzy assignments are identified and mapped along inferred trajectories to define regions of transition between cellular states (see *Identifying Fuzzy Cells*). Within these transition regions, genes associated with dynamic changes in expression are prioritised for downstream analyses, as described in *Identifying Major Regulatory Hubs*.

*Flufftail* then supports two complementary strategies to characterise gene regulatory network (GRN) dynamics. A data-driven strategy identifies candidate major regulatory hubs through a multi-step pipeline that integrates fuzzy gene module clustering with regulatory network inference (see *Identifying Major Regulatory Hubs*); a second pathway-guided strategy focuses on predefined gene sets of interest (see *Pathway-Guided Analysis*). For both approaches, regulatory co-variation is inferred and compared across stages to characterise state-dependent rewiring as described in *GRN Dynamics Across Pseudotime*).

### Fuzzy & Consensus Clustering

Cells in single-cell-resolution datasets often span a continuum of states, rather than forming strictly discrete groups. To capture this biological heterogeneity, we developed a fuzzy clustering framework based on repeated stochastic community detection. Clustering is repeated, across multiple iterations, using ClustAssess [9], with each run initialized using a different random seed, resulting in a set of *M* distinct partitions (Figure 1A). To ensure consistency of cluster-labels across partitions, we consolidated labels using the Hungarian algorithm [23]. The most frequently observed partition is selected as a reference; each subsequent partition is aligned to it by maximizing cluster overlap. This reconciled set of partitions provides the basis for computing both membership degree and consensus matrices.

Fuzzy clustering allows each cell to exhibit partial membership across multiple clusters. For each cell *x*_*i*_, we define its degree of membership *u*_*ij*_ in cluster *j* as the proportion of clustering iterations in which it is assigned to that cluster:

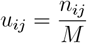

where *n*_*ij*_ is the number of times cell *x*_*i*_ is assigned to cluster *j*. The full membership matrix is given by *U* = [*u*_*ij*_]_*N* × *C*_, where *N* is the number of cells and *C* is the number of clusters; this matrix provides a probabilistic view of the clustering structure.

For downstream analyses requiring discrete labels, we also compute a “hard” (crisp) cluster assignment for each cell driven by the highest-probability cluster i.e. the cluster which it has the highest membership degree:

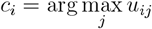

To quantify co-clustering consistency between cells, we compute a consensus matrix *S* = [*s*_*ij*_]_*N* × *N*_, where each entry reflects the proportion of clustering runs in which a given cell pair was co-assigned to the same cluster. Specifically, for each pair of cells *x*_*i*_ and *x*_*j*_, we define:

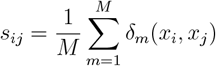

where *δ*_*m*_(*x*_*i*_, *x*_*j*_) = 1 if *x*_*i*_ and *x*_*j*_ are assigned to the same cluster in run *m*, and 0 otherwise. Further details are provided in Supplementary Method 1.

### Identifying Fuzzy Cells

*Flufftail* provides three complementary strategies to identify fuzzy cells: (i) low element-centric consistency (ECC), (ii) high entropy of cluster membership, and (iii) high entropy of co-clustering probabilities from the consensus matrix.

Cells with low ECC exhibit inconsistent cluster assignments across iterations (Figure 1A). ECC (also referred to as frustration) was introduced by Gates et al. [13] and is defined as:

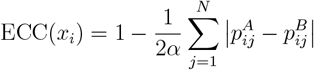

where *p*_*ij*_ is the personalized PageRank vector from node *i* to *j*.

Shannon entropy of the membership degree vector *u*_*i*_ quantifies the uncertainty in a cell’s cluster assignment. High entropy indicates diffuse membership across clusters:

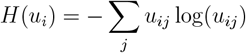

We additionally introduce an entropy-based metric, computed from the consensus matrix. Here, *p*_*ij*_ represents the probability that cells *i* and *j* are assigned to the same cluster across clustering iterations. Cells with high entropy have variable co-association patterns:

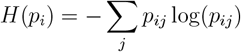

Further details and experiments are presented in Supplementary Method 2.

### Identifying Major Regulatory hubs

To identify key transcriptional drivers of transitional states (i.e. major regulatory hubs [24]), we developed a data-driven method to prioritize genes based on both expression fuzziness and regulatory influence across the inferred pseudotime. Starting with overlapping clusters across the pseudotime, i.e., clusters with high probability to transition from one to the other (Figure 1B), we define a *transition zone* by selecting cells between the centroids of the selected clusters, subsetting the pseudotime onto these cells, and identifying differentially expressed (DE), discriminative genes using Moran’s I spatial autocorrelation test (Figure 1C).

These DE genes are subjected to *fuzzy gene module clustering* using repeated stochastic community detection i.e. the fuzzy approach described earlier is applied on genes, rather than cells. This identifies “fuzzy genes” with low ECC, indicating co-membership across multiple modules. We hypothesise that genes participating in multiple modules, and thus active in diverse regulatory contexts, have an increased likelihood of being cross-modal regulators during cell state transitions. When such genes also exhibit high connectivity in GRNs, they become prime candidates for **major regulatory hubs** (perturbing them may propagate significant changes across modules and shift the global regulatory landscape).

To assess regulatory influence, we construct GRNs using GRNBoost2 [25], using each fuzzy gene as a candidate major regulatory hub and all DE genes as potential targets. For each major-regulatory-hub, we quantify its influence by calculating the proportion of targets for which its importance score exceeds the target-specific median and third quartile of importance values across all putative hubs. Candidate regulatory hubs are ranked based on this influence metric, and top-ranked genes are further analysed for GRN rewiring across pseudotime bins. All GRNs are inferred using the `reticulate` interface to GRNBoost2 from the `arboreto` package [25]. Further algorithmic and technical details are provided in Supplementary Method 3.

### Pathway-Guided Analysis

In addition to the data-driven strategy for identifying major regulatory hubs, *Flufftail* supports a pathway-guided analysis to accommodate prior biological knowledge. This strategy is intended for the targeted investigation of specific signalling pathways or functional gene sets relevant to the given biological context.

Following the identification of cell-state transitions, a gene set enrichment analysis using gprofiler2 [26] is performed to prioritise candidate pathways. For each pathway, annotated genes are intersected with the expression matrix, and used as input for downstream regulatory analysis within the *Flufftail* framework. This pathway-guided approach complements the unbiased data-driven strategy by enabling focused interrogation of predefined biological programmes.

Technical details of pathway selection, gene set handling, and implementation are provided in Supplementary Method 4.

### GRN Dynamics Across Pseudotime

Following the identification of candidate regulators using either the data-driven or pathway-guided strategy, we characterize the temporal (order-driven) dynamics of their regulatory interactions across the pseudotime trajectory. We infer GRNs at three discrete points (regulatory snapshots) along the transition (i) within the source cluster (cluster A), (ii) within the destination cluster (cluster B), and (iii) within the intermediate transition zone between them (Fig. 1C).

Each GRN is inferred using GRNBoost2, with expression matrices subsetted on the DE genes identified for the specified the transition. Top-ranked fuzzy genes, identified as candidate hub regulators, and all DE genes are treated as potential targets. To assess regulatory rewiring across pseudotime, we compare the topology and edge strength of the resulting GRNs. Interactions are filtered using quantile-based thresholds on importance scores, and changes in edge presence or magnitude are used to identify dynamic regulatory behaviours. GRNs are visualized individually and in combination, highlighting shared and condition-specific interactions across the transition.

## Results and Discussion

A first case study highlighting *Flufftail* is the end-stage liver single-nuclei RNAseq dataset [10] showing how clustering uncertainty can be leveraged to study cellular transitions between hepatocytes and cholangiocytes and regulatory dynamics. Fuzzy clustering underlines low-stability cells at the interface between these cell types and enables a data-driven prioritisation of major regulatory hubs. We then extend this analysis using a pathway-guided strategy to further characterise regulatory rewiring during the same transition. Next, we apply *Flufftail* to a spatial transcriptomics dataset from multiple sclerosis lesions [21] to illustrate the flexibility and scalability of *Flufftail* and how its GRN-dynamics component can be used to compare regulatory programmes across predefined tissue niches.

Full analyses for both case studies are provided in the Supplementary Analyses file.

### Fuzzy clustering highlights low-stability/ bi-phenotypic cells at the hepatocyte–cholangiocyte interface

To demonstrate the data-driven hub discovery strategy, we applied *Flufftail* on an end-stage liver single-nuclei RNAseq dataset [10], focusing on 23,835 cells spanning the hepatocyte– cholangiocyte transition zone (Supplementary Analysis A - Fig. S1).

Repeated stochastic clustering revealed a highly stable global partitioning of the data (median *ECC* > 0.95 at k=2, over 50 iterations). Within this otherwise stable clustering, *Flufftail* identified a subset of low-stability/fuzzy cells concentrated at the interface between the hepatocyte and cholangiocyte clusters, corresponding to the transition zone that was manually delineated in the original study and associated with bi-phenotypic cells (Fig. 1A). To identify an ordering of transition between the two states (hepatocytes and cholangiocytes, respectively) we computed the Monocle pseudotime trajectory (Fig. 1B).

To characterise transcriptional changes associated with this transition we selected cells centred around the centroids of the two stable clusters and their midpoint along the manifold (Fig. 1C). Differential expression analysis within this transition zone recovered genes previously linked to liver function, stress responses, and disease progression(Fig. 1D, Supplementary Analysis A) [10].

To provide a regulatory context for the static list of DE genes, we applied fuzzy gene module clustering to the transition-associated genes. This analysis identified genes with unstable/fuzzy module membership across repeated clustering runs, reflecting participation in multiple co-expression programmes. Applying the *Flufftail* major-regulatory-hub prioritization strategy highlighted several of the top 25 inferred major regulatory hubs corresponded to factors independently reported by Gribben et al. [10], including *FGF13, KLF6, CREB5*, and *SERPINE1*, all transcriptionally upregulated in the bi-phenotypic hepatocyte–cholangiocyte cells [10]. These results represent an external, data-driven validation of biological conclusions, illustrating the predictive potential of the *Flufftail* framework.

Visualisation of the fuzzy gene modules revealed that these candidate regulators localise in close proximity within the gene-module embedding (Fig. 1E), consistent with shared or overlapping regulatory roles. Analysis of their GRN dynamics showed a state-dependent handover of regulatory control: while *SERPINE1* orchestrates cytoskeleton and extracellular matrix remodeling (e.g., *SPIRE1, PLOD2*) in hepatocytes, *KLF6* and *FGF13* emerge as dominant drivers only within the transition zone. Here, they functionally converge on *CREB5* to form a dense regulatory core, with *FGF13* acting as a central hub for network rewiring, a topology that stabilizes and persists into the cholangiocyte state (Fig. 2A) [27].

**Figure 2.**
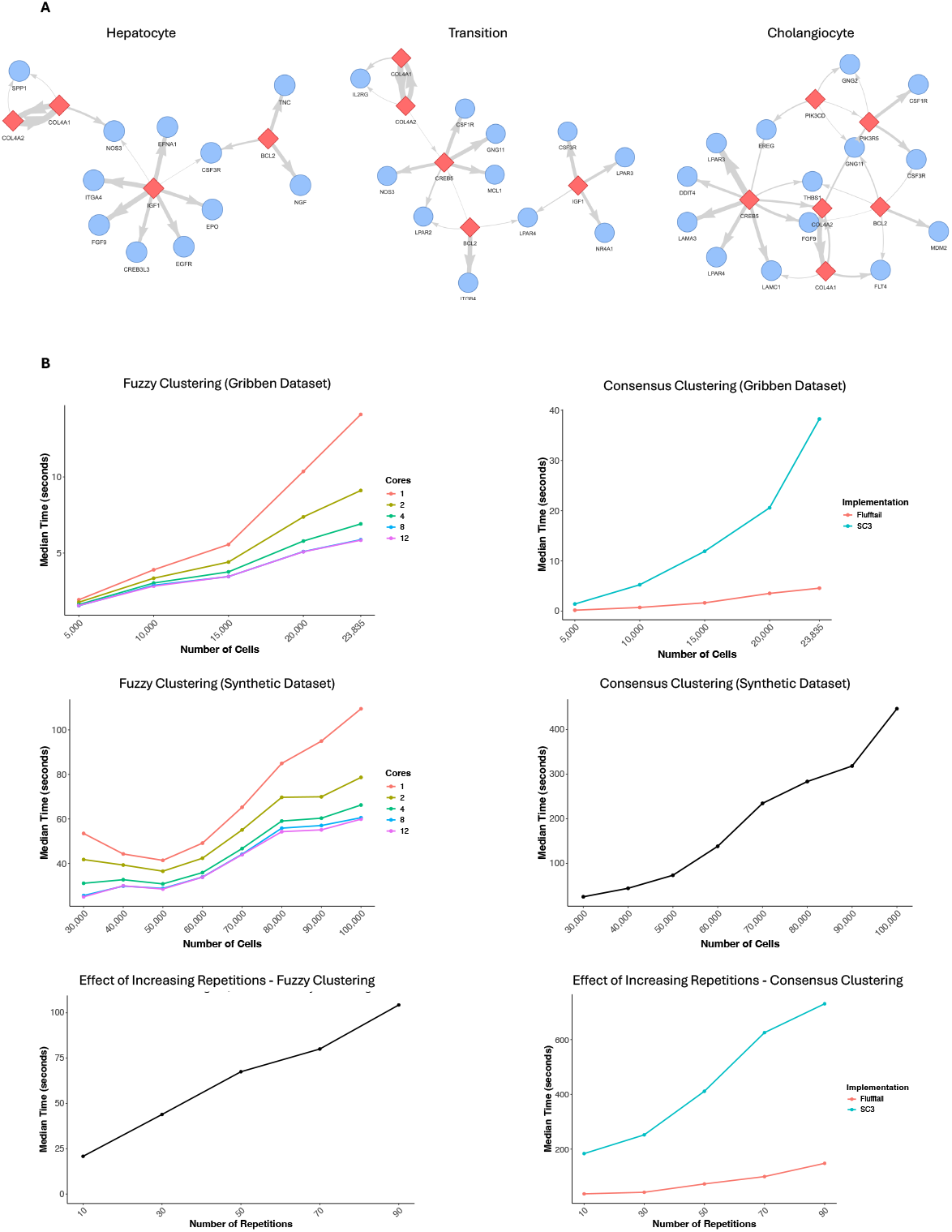
*Flufftail* reveals regulatory dynamics across lineage transitions and spatial niches. **(A)** Gene regulatory network (GRN) dynamics inferred for major regulatory hubs identified using the data-driven *Flufftail* strategy for the hepatocyte–cholangiocyte transition (*Gribben et al* study). Networks are shown for the hepatocyte state (LHS), intermediate transition state (middle), and cholangiocyte state (RHS), highlighting a state-dependent handover of regulatory control. *SERPINE1* dominates regulatory interactions in hepatocytes, whereas *KLF6* and *FGF13* emerge as key drivers within the transition zone, converging on *CREB5* to form a dense regulatory core that stabilises and persists into the cholangiocyte state. **(B)** GRNs inferred from spatial transcriptomics data of multiple sclerosis lesions across perilesional white matter, lesion rim, and lesion core (*Alsema et al* study). Using a curated set of disease-associated radial glia-like cell (DARG) and Sen-Mayo regulators, *Flufftail* captures niche-specific regulatory reorganisation, with *HMGB1, CDKN2A* (p16^Ink4a^), and *GDF15* exhibiting spatially distinct patterns of influence consistent with inflammatory and senescence-associated programs. **(C)** Screenshots of the *Flufftail* Shiny application, illustrating the interactive exploration of key analysis steps. The fuzzy clustering tab facilitates the inspection of hard cluster assignments, per-cell membership degrees, and consensus matrix computation, while the pseudotime tab assists with the trajectory inference and the identification of clusters with overlapping pseudotime distributions.

### Pathway-driven approach further characterises bi-phenotypic cells modulating hepatocyte-cholangiocyte dynamic

To complement the data-driven analysis, we used the pathway-guided strategy in *Flufftail* to examine regulatory dynamics within a predefined biological pathway, across cell states. We used the three cell populations defined using the centroid-based sampling described in the previous section, corresponding to the stable hepatocyte and cholangiocyte clusters, and the intermediate transition population, respectively.

Gene set enrichment analysis [28] was performed on the full expression matrix to prioritise pathways relevant to the dataset. This analysis highlighted the PI3K–AKT signalling cascade as significantly enriched(Fig. 1F) and consistent with findings from Gribben et al. [10]. Focusing on seven pathway-associated genes, including *IGF1, BCL2*, and *CREB5*, we used *Flufftail* to assess how their regulatory roles change across the centroid-defined cell states (Supplementary Fig. 2A, Supplementary Analysis A). *BCL2* is known to inhibit apoptosis; the proposed regulatory interactions underline this as an interesting factor, also associated with cancer; its involvement, in this context, confirms that hepatocytes transdifferentiation is associated with cellular stressed and potentially tumorigenesis.

In the hepatocyte state, *IGF1* and *BCL2* act as primary anchors. *BCL2* ‘s connections to *TNC* and *NGF* point to functions in extracellular matrix interactions and growth factor signalling, while *IGF1* coordinates broad associations with *EGFR, FGF9, EPO*, and *ITGA4*. A clear shift emerges in the transition zone, where *IGF1* redirects its influence toward alternative pathways, linking to *LPAR3, NR4A1*, and *CSF3R*. Notably, *ITGA4* undergoes a functional handover, disconnecting from *IGF1* and forming a strong interaction with *BCL2*, which in turn links the network to *CREB5*. By the cholangiocyte state, *IGF1* loses centrality, reflecting state-specific rewiring in which *CREB5* becomes the dominant regulator driving *LAMA3* and *EREG*. Together, these patterns illustrate how *Flufftail* can be used to interrogate dynamic regulatory rewiring within a specific signalling pathway across cell-state transitions.

### *Flufftail* reveals transcriptional dynamics on spatial transcriptomics datasets

To illustrate how the GRN-dynamics component of *Flufftail* can be applied to spatial data, we analysed a Visium spatial transcriptomics dataset of multiple sclerosis lesions from Alsema et al. [21]. Here, *Flufftail* was used to infer and compare GRNs across predefined spatial niches, rather than to identify cell states *de novo*. Spatial niches (perilesional white matter, lesion core, and rim) were used to subset cells for niche-specific GRN inference (Supplementary Analysis B).

Separate GRNs were inferred for each niche using the *Flufftail* GRN-dynamics workflow, focusing on transcriptional programmes driven by two curated sets of regulators: recently identified disease-associated radial glia-like cell (DARG) genes [29] and SenMayo genes [30]. These gene sets were used as candidate regulatory hubs, and regulatory interactions were inferred independently within each spatial compartment. Utilising a concatenation of these gene lists as the potential hubs revealed a reorganisation of GRNs across niches, with *HMGB1, CDKN2A*, and *GDF15* emerging as primary regulators (Fig. 2B).

Within the perilesional white matter, *HMGB1* exerted the largest influence on multiple targets (*NOTCH1, ICAM1, IL15*), all associated with generalised glial inflammation typical of this niche [21], with minimal regulation by *GDF15* and *CDKN2A*. Moving towards the rim of the lesion shifted the influence of the regulators: here, *GDF15, CDKN2A*, and *HMGB1* formed strong interactions with interferon-associated genes (*IFIT1, IFIT2, IFIT3, RFTN2, CHMP5*). In accordance with these findings, DARGs have been shown to be highly associated with senescence-related genes, including *HMGB1, CDKN2A*, and *GDF15*, within the lesion rim, where they induce IFN-associated inflammation in surrounding cells [31, 29].

The core of the lesion showed a further shift towards increased regulation by the senescence-associated genes *CDKN2A* and *GDF15*. These hubs were primarily associated with generalised inflammation (*ICAM1* and *GPR37*) as well as the IGF pathway (*IGFBP4*), with reduced involvement of *GDF15* as a regulatory hub. Recent work identified *CDKN2A* (p16^Ink4a^)-expressing senescent and inflammatory glial cells in the core of multiple sclerosis lesions [32, 33]. A complementary analysis using the differentially regulated gene set, identified by Alsema et. al [21], likewise showed that *Flufftail* recovered GRN changes consistent with pathway alterations across spatial compartments in multiple sclerosis tissue (Supplementary Analysis B).

### Robustness and scalability of the *Flufftail* framework

We benchmarked the *Flufftail* framework on the consensus/fuzzy clustering and consensus matrix components using both real and synthetic datasets (Supplementary Fig. 2B). For small to medium-scale benchmarks, we subsampled the Gribben et al. dataset [10] from 2,000 to 23,835 cells. On a personal computer (Apple M1 Max, 10 cores, 64 GB RAM), fuzzy clustering with 30 repetitions completed in approximately 10 seconds for datasets up to 20,000 cells using a single core, exhibiting near-linear scaling with sample size and substantial speedups with parallelization. Consensus matrix computation was 6–8 × faster than SC3 [16] across all *Gribben* subsets while producing identical outputs, owing to a C++/OpenMP optimization (Supplementary Method 1.3). To evaluate large-scale performance, we generated synthetic datasets of up to 100,000 cells with Splatter [22] and benchmarked on a 48-core server (754 GB RAM), confirming near-linear scaling for both fuzzy clustering and consensus computation. Runtime also scaled linearly with the number of clustering repetitions, with *R*^2^ = 0.992 for fuzzy clustering and *R*^2^ = 0.938 for consensus computation. In these experiments, *Flufftail* achieved 4–6× faster runtimes than SC3 across different numbers of repetitions.

### *Flufftail* enhances data mining tasks of regulatory interactions with an interactive interface

The *Flufftail* framework is accompanied by an interactive R Shiny interface that enables intuitive, end-to-end exploration of fuzzy clustering results and downstream regulatory analyses (Fig. 2C). The interface provides modular access to all key analytical steps, including inspection of membership degrees and element-centric consistency after fuzzy clustering, identification and visualisation of fuzzy cells and genes, pseudotime decomposition, differential expression analysis, fuzzy gene module clustering, and GRN inference. Through dynamic visualisations and parameter controls, users can interactively explore alternative clustering configurations, transition zones, and regulatory hubs without requiring extensive scripting, while retaining full reproducibility through the underlying R objects. This design facilitates rapid hypothesis generation, iterative exploration of biological signal versus technical variability, and transparent interpretation of clustering uncertainty, thereby extending the accessibility of *Flufftail* to researchers with diverse computational backgrounds and supporting data-driven investigation of cellular plasticity and GRN dynamics.

## Conclusions

In summary, the *Flufftail* framework provides a robust, reproducible, and scalable approach to uncovering regulatory dynamics in single-cell-resolution datasets, agnostic to modality, through fuzzy and consensus clustering. By systematically capturing clustering uncertainty, it delivers probabilistic, interpretable insights into changes/ dynamics in transcriptional signatures across cellular states.

The data-driven pipeline for identifying key regulatory hubs, integrated with GRN dynamics analyses across transitional stages, generates mechanistic insights into complex biological processes e.g. disease progression. In addition, the pathway-guided strategy allows targeted interrogation of regulatory rewiring within specific biological pathways, complementing the unbiased data-driven analysis.

Beyond regulatory dynamics, *Flufftail* provides a comprehensive framework for fuzzy clustering based on repeated community detection, an optimized implementation of consensus matrix construction, and robust ensemble partitions obtained by majority voting. While demonstrated here on single-cell and spatial transcriptomics data, the modular design of *Flufftail* makes it applicable to other high-dimensional omics modalities. For example, the framework could be directly applied to single-cell chromatin accessibility (scATAC-seq) or spatial metabolomics/proteomics datasets, where repeated clustering and centroid-defined state comparisons can be used to characterise dynamic changes in regulatory or functional influence.

## Supporting information

Main figure 2

Main figure 1

## Acknowledgements

We acknowledge the support of the Core Bioinformatics group; we thank Andi Munteanu and Floris Roos for early discussions during the development of this framework; we thank Eleanor Williams for constructive suggestions and assistance with reviewing the text. This research was funded by the Wellcome Trust [203151/Z/16/Z] and the UKRI Medical Research Council [*MC PC* 17230].

The Vallier lab is funded by core grants from the BIH and Max Planck Institute for Molecular Genetics, the ERC advanced grant Fun-Chol, and the Einstein Foundation.

For the purpose of open access, the authors applied a CC BY public copyright licence to all versions of the manuscript arising from this submission.

## Data Availability

The *flufftail* framework is freely available on GitHub at https://github.com/Core-Bioinformatics/ Flufftail and is accompanied by an interactive R Shiny interface that democratizes the analysis process, enabling researchers to leverage domain-specific expertise in extracting targeted hypotheses and meaningful insights from high-throughput data. The data used as case studies stemmed from publicly available datasets (*Gribben et al*, GSE202379, *Alsema et al*, GSE208747).

---

**Supplementary Figure 1.**
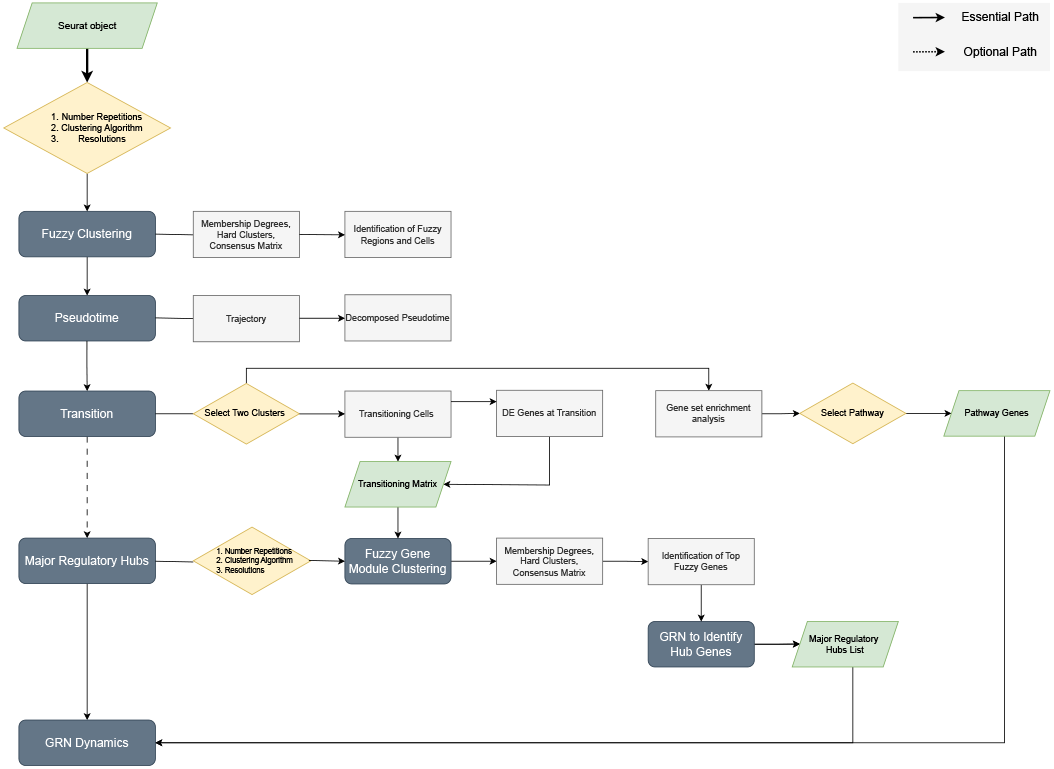
Workflow diagram summarising the *Flufftail* framework highlighting the essential and optional analytical paths. Starting from a Seurat object, repeated stochastic community detection is performed on a user-defined numbers of repetitions, clustering algorithms, and resolutions to derive fuzzy clustering outputs, including membership degrees, hard cluster assignments, and a consensus matrix. Element-centric consistency is used to identify fuzzy regions and cells, subsequently mapped along developmental trajectories using pseudotime inference. Transition zones are defined by selecting adjacent clusters with overlapping pseudotime distributions, from which transitioning cells and differentially expressed genes are identified. Two complementary strategies are then applied: a data-driven approach based on fuzzy gene module clustering of the transitioning matrix to prioritise major regulatory hubs via gene regulatory network (GRN) inference, and an optional pathway-guided approach based on gene set enrichment analysis. Both approaches converge on the inference of GRN dynamics to characterise regulatory rewiring across source, transition, and destination states.

**Supplementary Figure 2.**
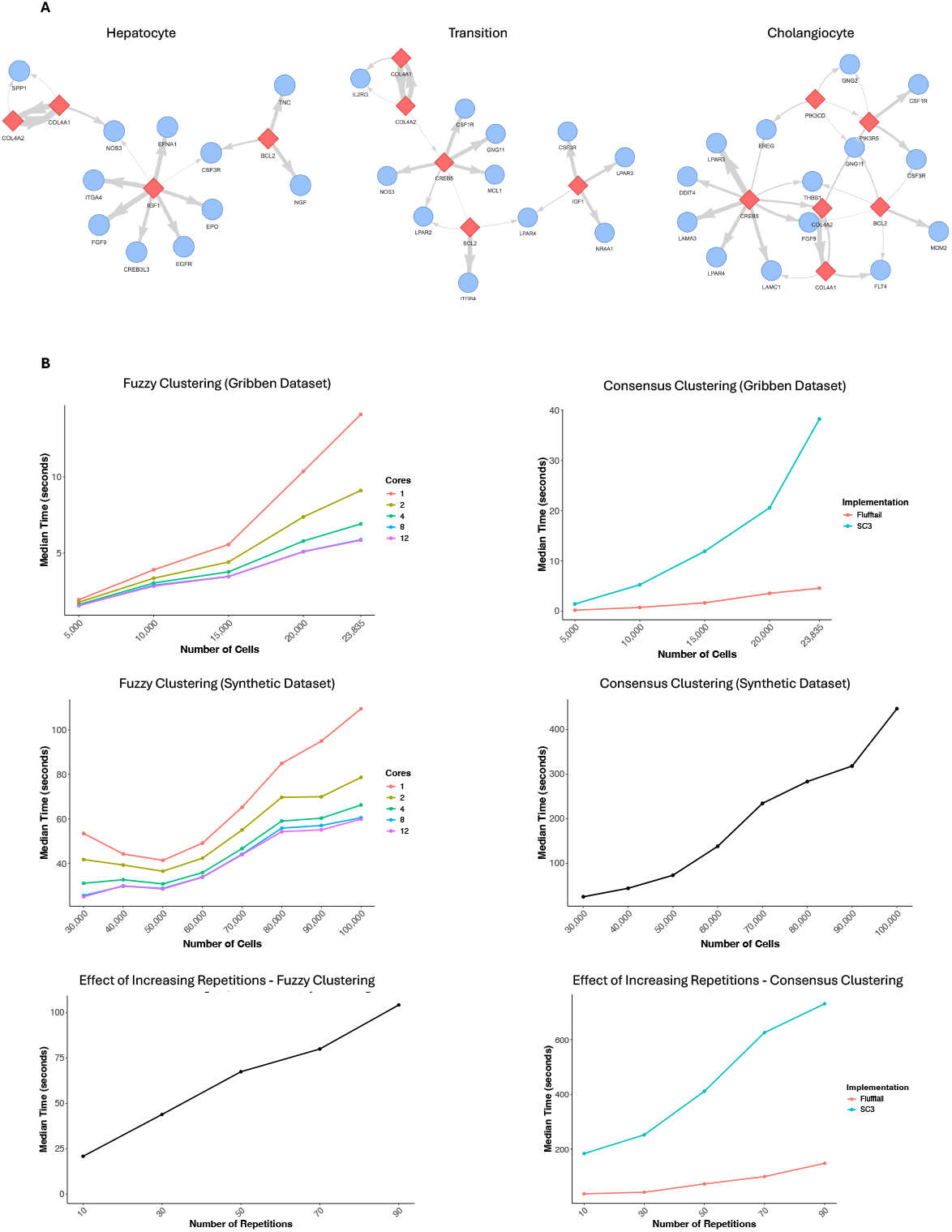
Pathway-guided GRN dynamics and scalability benchmarking. **(A)** Gene regulatory networks (GRNs) inferred using the pathway-guided *Flufftail* strategy for the PI3K–AKT signalling pathway (*Gribben et al* dataset), shown across three stages of the hepatocyte–cholangiocyte transition: hepatocyte state (LHS), intermediate transition state (middle), and cholangiocyte state (RHS). Networks highlight dynamic rewiring of regulatory interactions among key pathway-associated genes, including *IGF1, BCL2*, and *CREB5*, illustrating state-dependent shifts in centrality and target interactions during lineage progression. **(B)** Benchmarking of the *Flufftail* fuzzy clustering and consensus matrix components on real (Gribben et al.) and synthetic datasets. Median runtime is shown as a function of cell number, number of cores, and number of clustering repetitions, demonstrating near-linear scaling and substantially improved performance compared to SC3 across all evaluated settings.

