## Supplementary figures and images for "Unveiling Gene Regulatory Network Dynamics using Fuzzy Clustering"

### Main figure 1

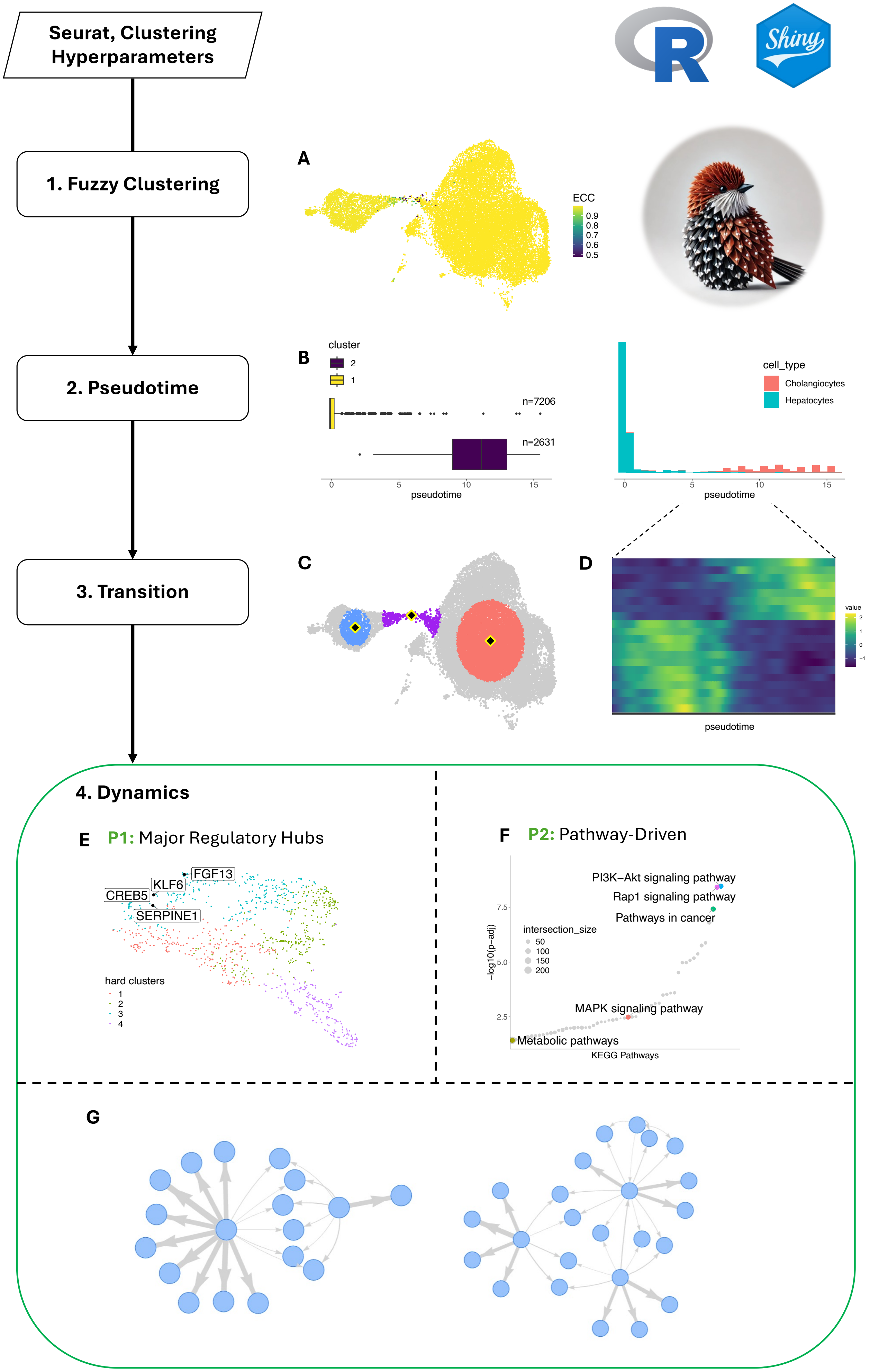

### Main figure 2

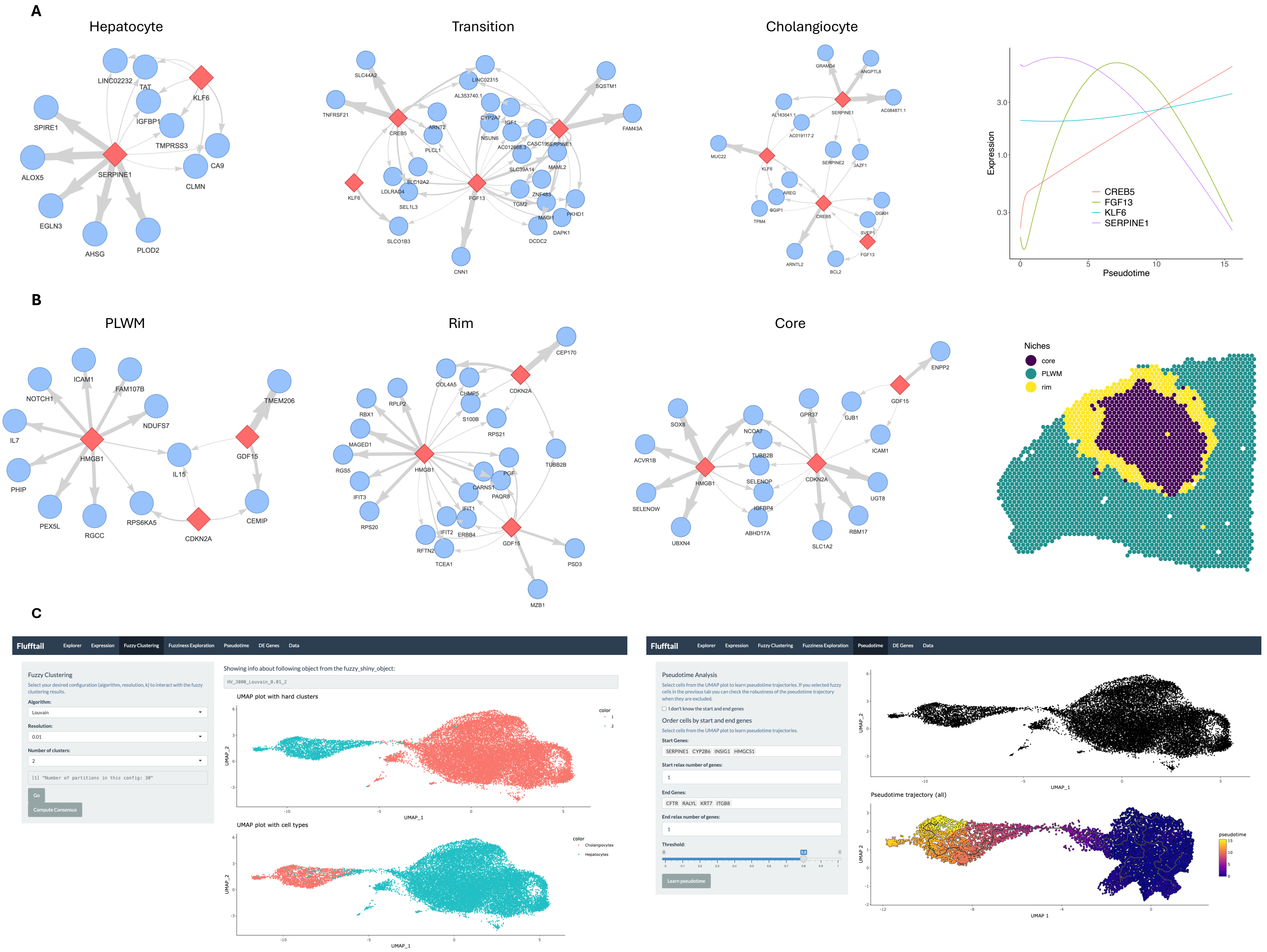
